# Predicting Transcriptional Responses to Unseen Compounds through Context Modeling and Multi-View Transfer

**DOI:** 10.64898/2026.11.05.764505

**Authors:** Yuqi Lei, Hanzhe Liang, Keqin Peng, Jun Wen

## Abstract

Predicting drug-induced transcriptional responses could support in silico screening of new compounds, yet existing models barely use the compound: on a drug-blind L1000 benchmark, removing the drug input from seven published models changes accuracy by at most 0.012 Pearson correlation. They capture shared cellular and experimental variation while overlooking the compound-specific signals that distinguish drug effects. We present UnDetect, a framework that separates contextual variation from compound effects to predict responses to transcriptionally unprofiled compounds. UnDetect integrates chemical structure, annotated targets, mechanisms of action and Cell Painting morphology into a normalized multi-view kernel that accommodates incomplete compound information, and transfers residual responses within and across profiled cell lines, so that pharmacological and phenotypic relationships in-form predictions beyond chemical similarity. On a drug-blind benchmark, where the paired control reveals the assay plate, UnDetect is the only model to beat a network that ignores the compound, in every fold; on a within-plate score that is zero for any compound-blind prediction, it reaches 0.117 against 0.092 for the strongest published model. On DeepCE’s benchmark it reaches 0.602 Pearson correlation against the reported 0.491. Transfer across cell lines improves 28 of 28 LINCS L1000 and, at the two higher doses, 47 and 49 of 50 Tahoe-100M cell-line evaluations. These results show that separating context from compound effects and integrating complementary compound information improves transcriptional prediction and extends the utility of perturbation atlases for virtual screening.

**Code:** https://github.com/CyclopsRay/UnDetect

## 1 Introduction

The transcriptional response of cells to a compound is a central readout in drug discovery: it reveals the compound’s mechanism of action, connects it to other compounds and to disease signatures, and suggests new uses for existing drugs [29,6]. Such profiling is now performed at atlas scale: the LINCS L1000 assay has measured 978 landmark genes after treatment with nearly 20,000 compounds [29], and Tahoe-100M has profiled 100 million single cells across 50 cancer lines [37]. Yet every new compound starts without any profile, and profiling each candidate across many cell lines is slow and costly. A model that predicts the response to an unseen compound, one never profiled, would turn these atlases into an in silico screen for new chemistry.

The task is hard for three reasons. First, the compound-specific part of a response is small compared with the cellular context, the cell line and the assay plate that all compounds measured together share. Second, chemical structure is a weak guide: a compound’s measured responses in other cell lines are a strong prior for a new line, whereas chemical fingerprints predict a held-out compound poorly [21,18]. Third, information about compounds is partial. Atlases are far from complete: in the drug-blind L1000 benchmark [3] a compound is profiled in 9 of 164 cell lines on average, almost always on a single plate per line, so less than 6% of the compound–line grid is measured; and annotated targets and Cell Painting morphology exist for only 36% and 23% of its compounds. Existing deep models encode the compound with a fingerprint or graph network [24,23,36,25,17], a pretrained molecular encoder [15,32,12] or a knowledge graph [10,12], and learn context and compound jointly by conditioning an expression decoder on that encoding.

These models barely use the compound. Bai et al. [3] retrained seven of them on a common drug-blind L1000 benchmark: replacing the drug input by zeros changes the accuracy by at most 0.012, and a multilayer perceptron that receives no drug information matches every one of them, as simple baselines do for genetic perturbations [1,35]. The reason is the plate. Each treated profile is paired with a control profile shared by all compounds on the same plate, so the control identifies the plate, and a plate-response baseline, which predicts a compound by the mean response of the other training compounds on its plate and uses no model and no drug information, outperforms five of the seven published models. Joint training trades one part for the other: the strongest published model, XPert, does learn the compound, as removing its compound input costs a third of its plate-centered score, but it falls below the drug-free network on the standard score.

Once the plate and cell line are taken into account, the response that remains is small, but it is shared across cell lines and predictable from how compounds relate to each other. We present UnDetect (UNseen Drug dEcomposition for Transcriptional rEsponse prediCTion), which fits the context first and the compound effect second (fig. 1). A backbone predicts the response that any compound would show in a sample. A multiview kernel over chemical structure, annotated targets, mechanism of action and Cell Painting morphology, normalized so that compounds lacking a view are not shrunk, generalizes the residual responses of profiled compounds to unprofiled ones, shared across cell lines and refined within each, and a bilinear interaction lets the cellular context rescale the compound effect gene by gene. Every step after the backbone has a closed-form solution.

**Fig. 1:**
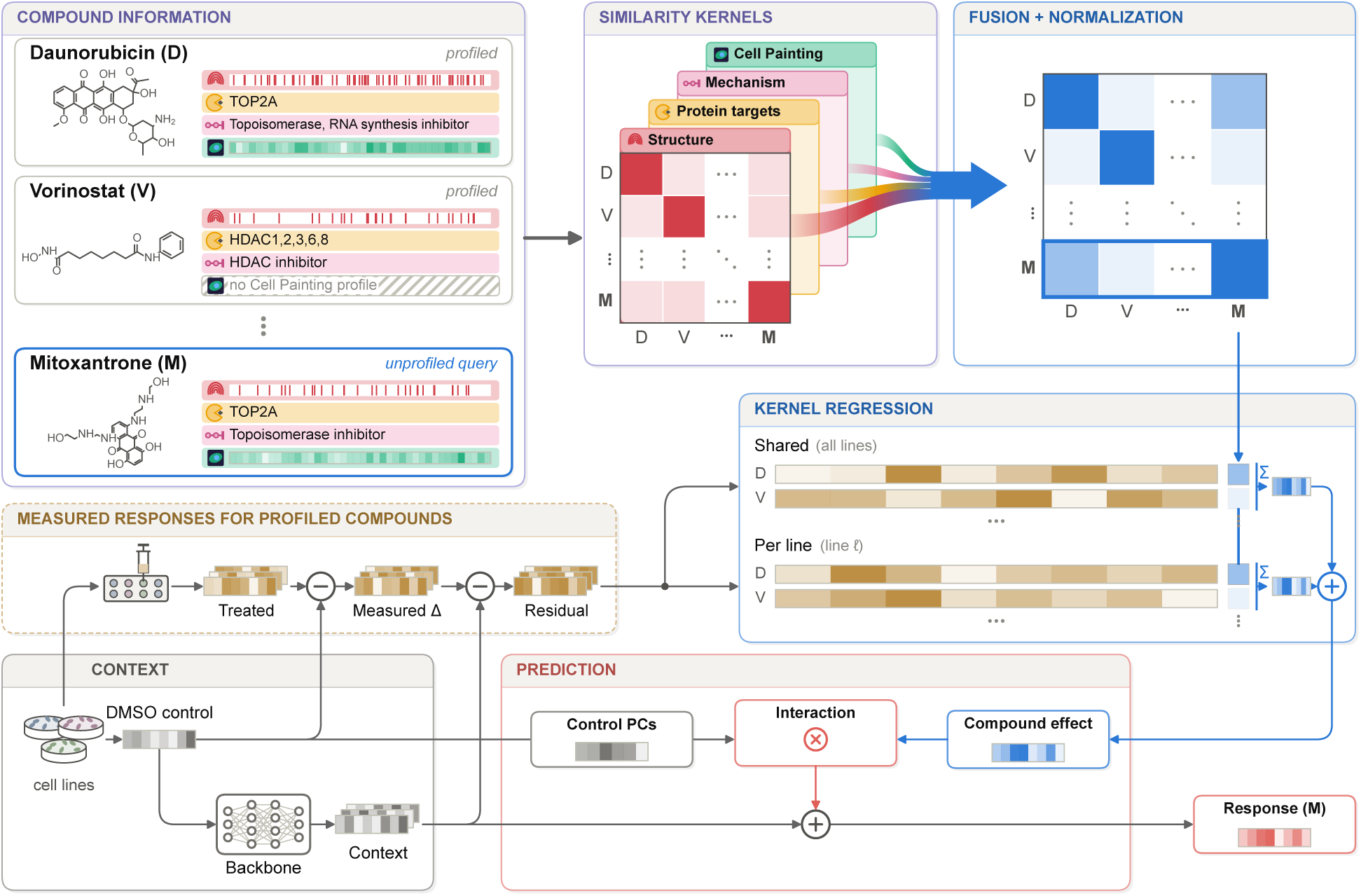
UnDetect, shown for two profiled compounds (daunorubicin, D; vorinostat, V) and an unprofiled query (mitoxantrone, M). *Compound information*: four views per compound, hatched where missing. *Similarity kernels*: one per view. *Fusion + normalization*: the views weighted by their alignment with training responses (ribbon widths); the fused kernel pairs M with D, which shares its target and mechanism, rather than with V, its nearest structure. *Context*: a backbone on the DMSO control predicts the response shared by all compounds; subtracted from the *measured responses*, it leaves the residuals of profiled compounds. *Kernel regression*: the query’s compound effect from these residuals, shared across lines and per line. *Prediction*: the control’s principal components rescale the compound effect gene by gene, and the context is added.

Our contributions are threefold. (i) A closed-form model for unseen compounds that fits the compound effect after the context, shares it across cell lines and lets the context rescale it gene by gene (section 2). (ii) A multi-view compound kernel normalized so that the effect a compound can receive does not shrink with the views it lacks (section 2.3). (iii) The finding that the paired control encodes the plate, and plate-centered scores that isolate the compound-specific response (section 3.1). On the drug-blind benchmark UnDetect is the only model above the drug-free baseline on the standard score, in every fold. On the within-plate score, which is zero for any prediction that ignores the compound, it raises the correlation of the strongest published model from 0.092 to 0.117, and it picks out a compound’s own measured signature among those of its plate in 27% of cases, where that model does so in 19%. For compounds known only by structure the two are level; UnDetect’s lead comes from the pharmacological and phenotypic views. On DeepCE’s benchmark it reaches a Pearson correlation of 0.602, where DeepCE reported 0.491 (section 3).

## 2 Method

### 2.1 Task definition

Each sample *i* is a compound *d_i_* applied to a cell line *ℓ_i_*, with response *y_i_ ∈* R*^G^* over the *G* = 978 L1000 landmark genes *j*. Where each treated profile *x*_trt_*_,i_* is paired with a control profile *x_i_*, measured on cells of the same line on the same plate treated with the vehicle (DMSO) alone, the response is *y_i_* = *x*_trt_*_,i_ − x_i_*; where the control has already been removed, *y_i_* is the given signature (section 3.1). *T* is the set of training samples, *D* the set of training compounds and *D_ℓ_ ⊆ D* those measured in line *ℓ*. The task is to predict *y_i_* for an unseen compound, one absent from training in every cell line (drug-blind); all cell lines are seen. We write the prediction as a context term, the response that any compound would show in that sample, plus a compound effect, and fit the two in turn (fig. 1):

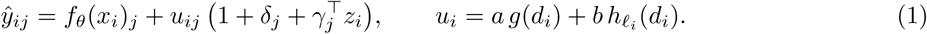

The backbone *f_θ_* gives the context (section 2.2); *g* and *h_ℓ_* are the shared and line-specific effects of a compound, from a multi-view compound kernel (sections 2.3 and 2.4); and *δ_j_*, *γ_j_* let the context *z_i_* rescale the compound effect on gene *j* (section 2.5). Every stage after the backbone has a closed form.

### 2.2 Context backbone

The backbone *f_θ_*sees no compound information, and its residuals

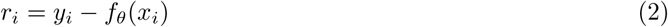

are what the compound effect must explain. It is the drug-free network of Bai et al. [3], a multilayer perceptron on the control profile (three hidden layers of 2,048 units), trained on training compounds only and then frozen; because the control identifies the plate, it absorbs cellular and plate context. Where the control has already been removed, the backbone is the training mean of each line. Fitting the backbone first keeps the large, compound-independent part of the response from swamping the compound part; jointly trained models trade one for the other (section 3.3). Supplementary Appendix A.1 gives the training details.

### 2.3 Multi-view compound kernel

Each view *v* gives a positive semidefinite kernel *K_v_* over all compounds, with unit diagonal on the compounds where the view is observed: Tanimoto similarity of Morgan fingerprints [26] (radius 2, 2,048 bits; all compounds), Tanimoto similarity of annotated target sets and of mechanism-of-action sets [6,37], and cosine similarity of *z*-scored Cell Painting profiles at 10 *µ*M [4,34,13]. A compound lacking view *v* gets a zero row and column in *K_v_*, which keeps *K_v_* positive semidefinite. The views are fused and normalized,

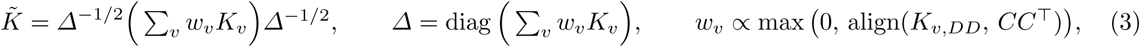

where *C* holds one row per training compound, its mean residual *r* over all its training samples, and the weight *w_v_* is the centered alignment [7] between *K_v_*on the training compounds and the Gram matrix of *C*, set to zero if negative and normalized to sum to one.

Zero-filling a missing view shrinks the compound toward the backbone. Let *S_d_* be the views observed for compound *d*. Then ^∑^*_v_ w_v_*(*K_v_*)*_dd_* = ∑*_v∈Sd_ w_v_*, and by the reproducing property every function *ϕ* in the reproducing kernel Hilbert space of ∑*_v_ w_v_K_v_* satisfies *|ϕ*(*d*)| ≤ |ϕ|(∑*_v∈Sd_ w*)^1/2^. Kernel ridge regression bounds |*ϕ*|, so the largest effect a compound can receive shrinks with the views it lacks, whatever its response. The normalization by *Δ* in eq. (3) is a congruence, so *K̃* stays positive semidefinite, and it gives every compound unit self-similarity: the views that are present set its similarities, and the bound is the same for every compound, whichever views it has. This removes the shrinkage that zero-filling imposes; how much of a compound’s effect can be predicted still depends on the views it has (table 5). Structure is observed for every compound, so *Δ* is invertible; Supplementary Appendix A.1 reports the view weights.

### 2.4 Generalization across and within cell lines

Kernel ridge regression with penalty *λ* extends the mean residuals to any compound *d*, as a shared and a line-specific effect,

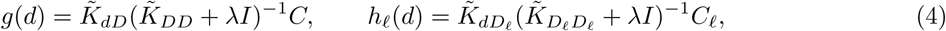

where *C_ℓ_* holds the mean residuals of the compounds of *D_ℓ_* in line *ℓ*. The shared effect *g* regresses the mean residuals of all training compounds, so a compound profiled in one cell line informs every cell line; the linespecific effect *h_ℓ_* uses only the compounds measured in line *ℓ*. Equivalently, *g* minimizes |*C − K̃*_*DD*_α|^2^ + *λ* tr(*α^⊤^K̃_DD_ α*) over coefficients *α*, and the scales *a* and *b* in eq. (1) set how much each term contributes. When doses are resolved, as in the DeepCE benchmark, both terms are fitted jointly as one kernel ridge regression over samples (*d, ℓ, s*), where *s* is the dose in *µ*M, with the coregionalized kernel [2]

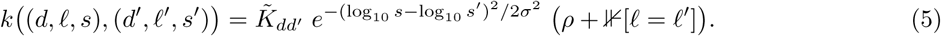

Here *σ* sets how fast similarity decays with dose and *ρ* how much of a compound’s effect is shared between cell lines; *ρ* = 0 recovers independent per-line models, and eq. (4) is the two-stage approximation with *a/b* in the role of *ρ*. Kernel models over compounds and cell lines are established in drug-response prediction, as multitask models [11,8] and as products of a compound and a cell-line kernel [5], which contain independent per-line models as a special case [28]; UnDetect uses the normalized multi-view *K̃* as the compound kernel.

### 2.5 Compound–context interaction

*Role.* The kernel terms give a compound the same effect profile in every sample of a cell line, up to *a* and *b*, but cells in a different state can respond to the same compound more or less strongly in particular genes. In eq. (1), 1 + *δ_j_* + *γ^⊤^z_i_* rescales the compound effect on gene *j*: *δ_j_* in every sample, *γ_j_* in proportion to the context. This is a varying-coefficient model [14], akin to feature-wise linear modulation [22].

*Context. z_i_* holds the scores of *x_i_* on the leading *q* principal components of the distinct training control profiles, standardized. It is defined for any sample with a control profile.

*Closed form.* The coefficients *β_j_* = (*δ_j_, γ_j_*) minimize the penalized squared error

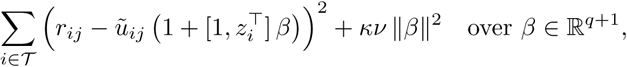

which gives one (*q*+1) *×* (*q*+1) linear system per gene,

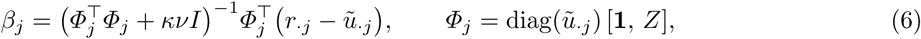

with the contexts *z_i_* stacked in *Z*. The factor 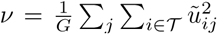 puts the penalty *κ* on the scale of the compound effect. At *κ* = *∞*, *β_j_* = 0 and eq. (1) reduces to the kernel terms, so model selection can always switch the interaction off.

*Leave-one-compound-out inputs.* Fitted on in-sample compound effects, the interaction would learn to undo the shrinkage of the kernel regression. *ũ_i_* = *a g̃*(*d_i_*) + *b h̃ℓ* (*d_i_*) therefore uses the value of each term with compound *d_i_* left out, which kernel ridge regression gives in closed form,

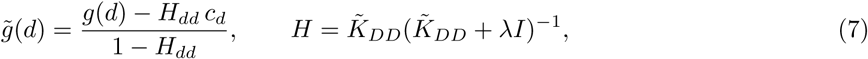

with *c_d_* the row of *C* for *d*, and the same within each line for *h̃ℓ*. This holds the view weights fixed; compoundstage cross-fitting that re-estimates them gives the same gain (Supplementary Table 15).

### Without a per-sample control

On LINCS the context is the cell line: 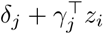 is replaced by one scale *δ_ℓj_*per cell line and gene, fitted by the same ridge regression on the cross-validated compound effects of the training compounds, with *κ* chosen by cross-fitting over the inner folds.

### Related designs

Published models couple context and compound inside one network (section 3.2), also through a nonlinear decoder over additive latent embeddings [19,15]. Tensor completion over compound, gene and cell line needs every compound profiled in some cell line [16], and response decompositions [21] and observed-statistics models, which express a response through the target line’s control profile (Rhaister-O) or through a small panel of drugs profiled in it (Rhaister) [30], carry a compound’s measured responses to new cell lines; for an unseen compound, UnDetect predicts these compound terms from its views.

### Model selection

The ridge *λ*, the scales *a*, *b* and *κ*, the number *q* of components (and *ρ*, *σ* when doses are resolved) are chosen from fixed grids by the benchmark’s own score on validation compounds, which are disjoint from training and test compounds, or by compound-grouped inner cross-validation on LINCS and Tahoe-100M; each test set is scored once. All compound views are fixed annotations or measurements in other assays; no response of a test compound is used at any stage. Supplementary Table 6 lists the grids.

### Computational cost

After the backbone, UnDetect solves linear systems only; one eigendecomposition of *K̃_DD_* serves every *λ* and every leave-one-compound-out value. On one A100 GPU, fitting UnDetect on a fold of the drug-blind benchmark takes about 5 minutes (backbone and kernel terms 3.2 minutes, interaction 2 minutes, 5 GB of GPU memory), against 268 minutes for training XPert on the same fold.

## 3 Experiments

### 3.1 Datasets and setup

We use four unseen-compound benchmarks (table 1). On the drug-blind benchmark of Bai et al. [3] the response is the treated profile minus its paired control, and the metric is the mean over test samples of the Pearson correlation between true and predicted responses across all 978 genes (PCC_DEG_, named as in [3]). The other three use plate-normalized Level-5 signatures [29], or pseudobulk deltas for Tahoe-100M, and report the Pearson correlation per signature or per (compound, line), averaged over lines. The drug-blind and LINCS benchmarks have one dose and time point (10 *µ*M, 24 h); Tahoe-100M is scored separately at each of its three doses, and only DeepCE’s benchmark resolves doses within one model. Supplementary Table 21 lists the data sources and versions.

**Table 1:**
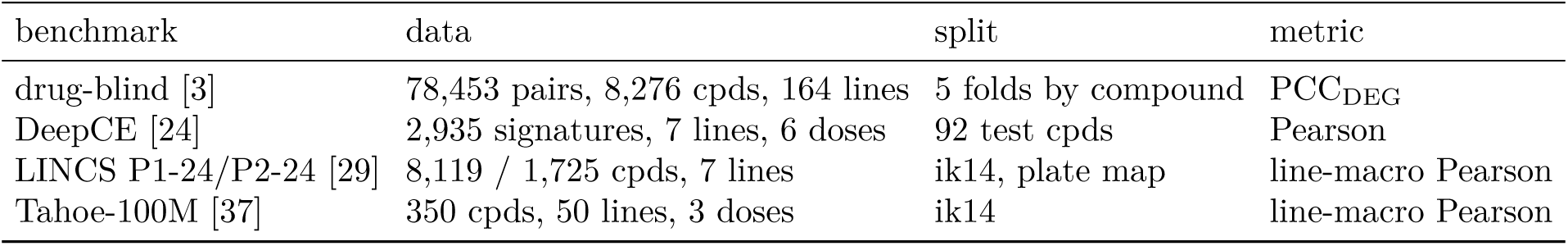
Benchmarks. All hold out whole compounds (drug-blind: train, validation and test 3:1:1 per fold). P1-24 / P2-24: Phase I / II, 24 h. DeepCE: its published split. ik14: compounds grouped by the first InChIKey b<u>lock; plate map: compounds sharing an L1000 plate map kept together.</u>

| benchmark | data | split | metric |
|---|---|---|---|
| drug-blind [3] | 78,453 pairs, 8,276 cpds, 164 lines | 5 folds by compound | PCC<sub>DEG</sub> |
| DeepCE [24] | 2,935 signatures, 7 lines, 6 doses | 92 test cpds | Pearson |
| LINCS P1-24/P2-24 [29] | 8,119 / 1,725 cpds, 7 lines | ik14, plate map | line-macro Pearson |
| Tahoe-100M [37] | 350 cpds, 50 lines, 3 doses | ik14 | line-macro Pearson |

*Plate structure.* In the drug-blind benchmark, the control takes 9,951 distinct values, exactly one per pair of cell line and detection-plate set: every treated sample on a plate shares the same control vector, up to 321 samples per control. Because the split holds out compounds rather than plates, 90% of test samples share their control, and hence their plate, with training samples, so a model that receives the control can recognize the plate. The other three benchmarks remove the shared plate response before modeling (robust *z*-scores against the population of the same plate [29], or subtraction of the plate’s DMSO profile); on LINCS we also hold out compounds together with their plate maps.

### Plate-centered metrics

Plate structure inflates PCC_DEG_ for any model that can see the control, and a prediction of the plate mean is of no use for a compound that has not yet been assayed. We therefore also score the deviation of a compound from the other compounds on its plate. For a test sample *i* with training plate-mates, let *m_i_* be their mean response; the plate-centered score is

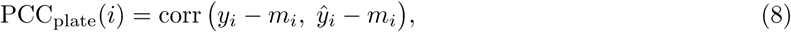

the correlation across genes, averaged over test samples. The error of *m_i_* enters both terms, so a compoundblind prediction can score above zero. The within-plate score centers truth and prediction each on its own mean over the test compounds of a plate *p*, PCC_within_(*p, d*) = corr(*y_pd_ − y̅_p_, y*^*_pd_ − y̅*^*_p_*), averaged over (plate, compound) pairs. Any prediction constant within a plate, and hence any prediction that ignores the compound, scores zero. PCC_within_ is our main compound-specific score and PCC_plate_, covering more samples, its complement; all our settings are chosen by PCC_DEG_, so neither is optimized. Samples without a training plate-mate are scored with PCC_DEG_ and reported separately. This is the chemical-perturbation counterpart of centering on systematic variation in genetic perturbation benchmarks [33]; here the systematic component has a concrete source, the shared control. Supplementary Appendix A.11 defines PCC_within_ in full, and Table 18 gives it per fold.

### 3.2 Compared methods

On the drug-blind benchmark we compare with the seven published models retrained by Bai et al. [3], CIGER [23], MultiDCP [36], PRnet [25], PertDiT [17], DeepCE [24], TranSiGen [32] and XPert [12], scored from their released predictions, and with their drug-free network, a multilayer perceptron on the control alone. The three strongest of them couple the compound and the context in different ways. DeepCE [24] encodes the molecular graph with a graph convolutional network and relates its substructures to genes by multi-head attention. TranSiGen [32] learns latent spaces of the control and treated profiles with variational autoencoders and maps the control latent, together with a pretrained molecular embedding, to the treated one. XPert [12] turns the control profile into gene tokens that attend to the atoms of a pretrained three-dimensional molecular encoder, and adds a compound embedding pretrained on a graph of protein interactions, drug–target links and drug–drug similarities. The *plate-response baseline* predicts a test sample by the mean response of the training samples that share its control; it uses no compound information and fits no parameters. The *reference* predicts each test sample by the backbone plus the mean residual of the same compound’s other test samples (other cell lines); it uses test responses and marks how much a compound’s own measurements add. On DeepCE’s benchmark we compare with the models reported by Pham et al. [24], our reproduction of DeepCE, a tuned Morgan nearest-neighbor search, and a neural network given the same compound views as UnDetect. On LINCS and Tahoe-100M we compare with the training mean of each line and with per-line kernel regressions on chemistry alone or on all views.

### 3.3 Generalization to unseen compounds

On the drug-blind benchmark, no published model beats a network that ignores the compound on PCC_DEG_, and UnDetect does in every fold (table 2, fig. 2a). The plate-response baseline alone, which looks up the mean response of a sample’s plate-mates, is above five of the seven published models; on the samples that have a training plate-mate it scores 0.643, above all seven, and on the others it falls to the level of the cell-line mean. The strongest published model, XPert, stays below the drug-free network, and UnDetect is above it in every fold with bootstrap intervals above zero. Supplementary Tables 7 and 8 give the scores per fold.

**Fig. 2:**
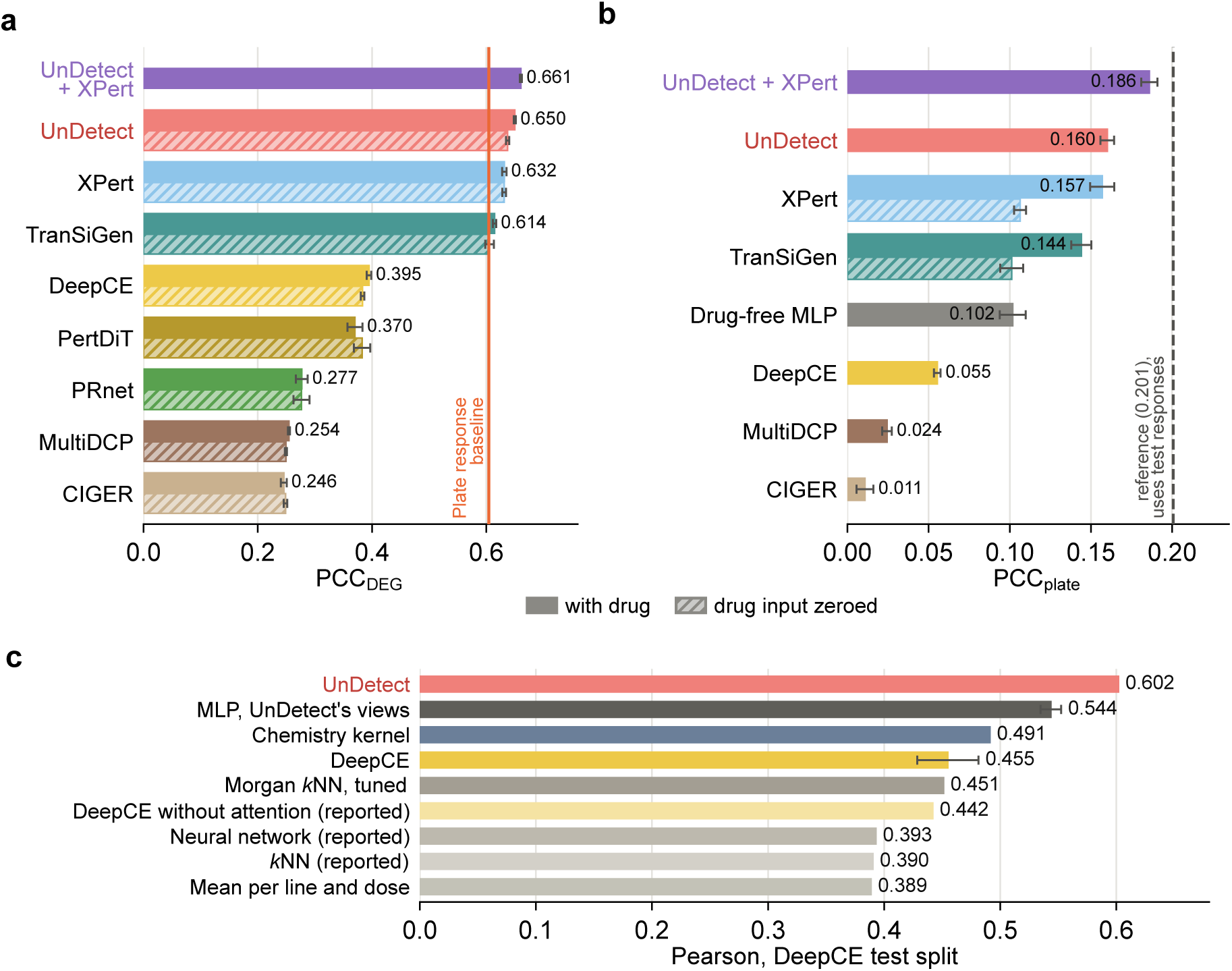
Unseen-compound generalization (a, b: drug-blind benchmark, mean of five folds, error bars: s.d. over folds; c: DeepCE’s benchmark, one published split). (a) PCC_DEG_ of the published models with their drug input (solid) and with it replaced by zeros (hatched) [3], and of UnDetect (hatched: its drug-free backbone); line: the plate-response baseline. (b) PCC_plate_ on the 90% of test samples with a training plate-mate; dashed: the reference, which uses the compound’s measured responses in other cell lines. (c) Per-signature Pearson (error bars: s.d. over training seeds where several were run); DeepCE is our reproduction; (reported): as reported by Pham et al. [24]; MLP: our network given the same compound views as UnDetect.

**Table 2:** Drug-blind benchmark, mean of five folds. No mate: PCC_DEG_ on the 10% of test samples without a training plate-mate; PCC_plate_ on the other 90%. *Δ*: difference to the drug-free MLP, which our backbone reproduces (0.637, 0.100). Bold: best; underlined: best single model. Published models: predictions released by Bai et al. [3]; PRnet and PertDiT are released on another scale, so only their PCC_DEG_ is shown. Ensemble: 0.6 UnDetect + 0.4 XPert, the weight chosen on another fold’s test compounds (0.4 in every fold); a fixed weight of 0.5 is also above both models in every fold. Reference: uses test responses (section 3.2).

| model | PCC<sub>DEG</sub> | *Δ* | no mate | PCC<sub>plate</sub> | *Δ* | PCC<sub>within</sub> |
|---|---|---|---|---|---|---|
| plate-response baseline | 0.604 | −0.033 | 0.239 | 0 | −0.102 | 0 |
| CIGER | 0.246 | −0.392 | 0.260 | 0.011 | −0.091 | – |
| MultiDCP | 0.254 | −0.383 | 0.224 | 0.024 | −0.077 | – |
| PRnet | 0.277 | −0.360 | – | – | – | – |
| PertDiT | 0.370 | −0.267 | – | – | – | – |
| DeepCE | 0.395 | −0.243 | 0.338 | 0.055 | −0.046 | 0.027 |
| TranSiGen | 0.614 | −0.023 | 0.564 | 0.144 | +0.042 | 0.079 |
| XPert (best published) | 0.632 | −0.006 | 0.565 | 0.157 | +0.055 | 0.092 |
| drug-free MLP | 0.637 | – | 0.535 | 0.102 | – | 0 |
| UnDetect | 0.650 | +0.013 | 0.574 | 0.160 | +0.058 | 0.117 |
| UnDetect + XPert (ensemble) | **0.661** | **+0.023** | **0.614** | **0.186** | **+0.084** | **0.126** |
| reference (test responses) | – | – | – | 0.201 | +0.099 | – |

The gain is compound-specific. On PCC_within_ UnDetect is the best single model in every fold (table 2). On PCC_plate_ it raises the backbone’s score by more than half, in every fold (fig. 2b); the backbone scores above zero there because each plate mean rests on finitely many plate-mates. Without any measurement of the compound, UnDetect attains 60% of the gain of the reference, which uses the compound’s own measurements in other cell lines. On test samples with no training plate-mate, where the plate response cannot be looked up, UnDetect is also ahead of both the drug-free network and XPert. Supplementary Figure 5 and Table 17 show PCC_plate_ by the number of plate-mates.

The plate-centered scores show that XPert learns the compound at the cost of the context. It is level with UnDetect on PCC_plate_ and loses a third of that score when its compound input is replaced by zeros, although its PCC_DEG_ does not change; TranSiGen shows the same pattern. Yet XPert stays below the drugfree network on PCC_DEG_, whereas UnDetect, which fits the compound after the context, gains on both scores and leads XPert on PCC_within_, 0.117 against 0.092, in every fold with intervals above zero. A screen needs predictions that tell compounds apart: among the test compounds of a plate, UnDetect’s prediction for a compound is closest, in correlation across genes, to that compound’s own measured signature in 27% of cases, against 19% for XPert and 11% by chance (one over the number of test compounds on the plate, averaged), in every fold. The two models capture different parts of the compound signal: their weighted average is above both on PCC_DEG_ and PCC_plate_ in every fold, also with a fixed weight. Supplementary Table 12 gives every published model with and without its drug input, and Table 19 the retrieval by plate size.

### DeepCE benchmark

On DeepCE’s own benchmark, UnDetect reaches 0.602 per-signature Pearson against 0.491 reported for DeepCE (fig. 2c), and improves 79 of the 92 test compounds over our reproduction of DeepCE; a neural network given exactly the same inputs and tuned on the same validation set stays well below it. Supplementary Table 9 lists every model on this benchmark.

*LINCS and Tahoe-100M.* Here the backbone is the training mean of each cell line, so the whole compound effect comes from the kernel. UnDetect is best in every setting (table 3): over the per-line multi-view kernel, sharing the compound effect across cell lines improves every LINCS line in all four settings and nearly every Tahoe-100M line at the two higher doses. The shared term carries compound-specific information to lines that have not profiled the compound, and its gain is largest when each training compound is profiled in few lines, as in the separation of shared and line-specific response components in genetic screens [21]. Supplementary Figure 4a shows this gain by coverage.

**Table 3:**
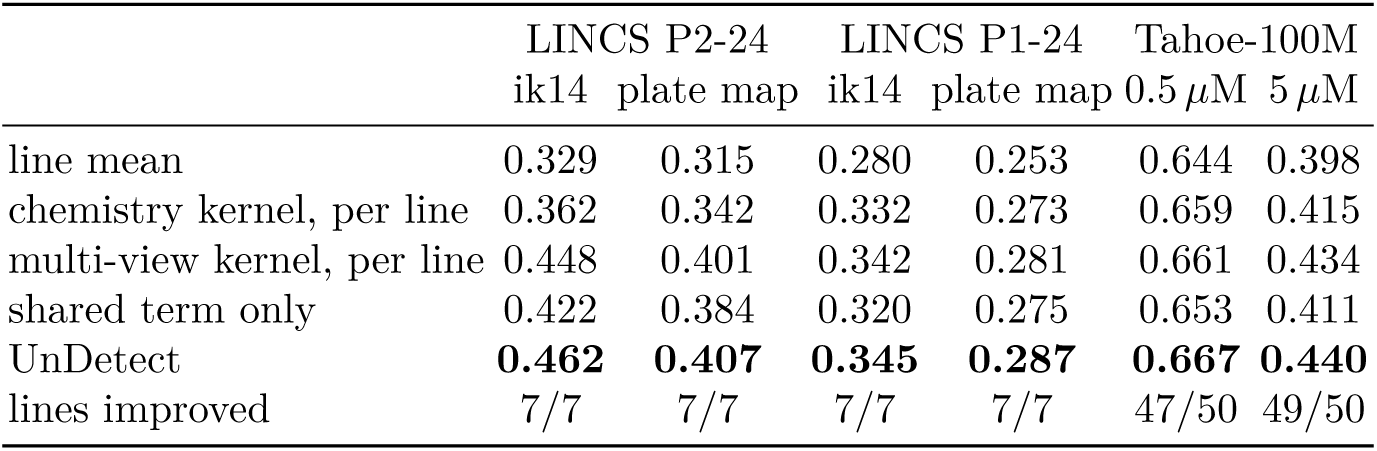
LINCS and Tahoe-100M, line-macro Pearson (unseen compounds). Per line: one kernel model per cell line; Shared term only: the cross-line term of UnDetect alone. UnDetect: per-line plus shared term, without the interaction. Lines improved: UnDetect over the per-line multi-view kernel. At 0.05 *µ*M every model is within <u>0.011 of the line mean.</u>

|  | LINCS P2-24 |  | LINCS P1-24 |  | Tahoe-100M |  |
|---|---|---|---|---|---|---|
|  | ik14 | plate map | ik14 | plate map | 0.5 *µ*M | 5 *µ*M |
| line mean | 0.329 | 0.315 | 0.280 | 0.253 | 0.644 | 0.398 |
| chemistry kernel, per line | 0.362 | 0.342 | 0.332 | 0.273 | 0.659 | 0.415 |
| multi-view kernel, per line | 0.448 | 0.401 | 0.342 | 0.281 | 0.661 | 0.434 |
| shared term only | 0.422 | 0.384 | 0.320 | 0.275 | 0.653 | 0.411 |
| UnDetect | **0.462** | **0.407** | **0.345** | **0.287** | **0.667** | **0.440** |
| lines improved | 7/7 | 7/7 | 7/7 | 7/7 | 47/50 | 49/50 |

### 3.4 Ablation study

Each component of UnDetect raises the compound-specific score (table 4): the chemistry kernel gives about half of the gain, the multi-view kernel most of the rest, and the interaction improves both metrics further, in every fold with intervals above zero. A kernel over plate co-membership, which knows which compounds were assayed together and nothing about them, adds nothing on PCC_plate_ and PCC_within_; its PCC_DEG_ gain, half of UnDetect’s, comes from the plate alone, which is why we measure compound learning on the platecentered scores. Rescaling by the identity of the cell line instead of its control profile gives a similar gain on this benchmark, where every line is seen in training; we use the control profile because it is also defined for a line never profiled. Supplementary Table 15 shows that permuting all compound views jointly removes the gain, and Table 11 compares the interaction variants.

**Table 4:**
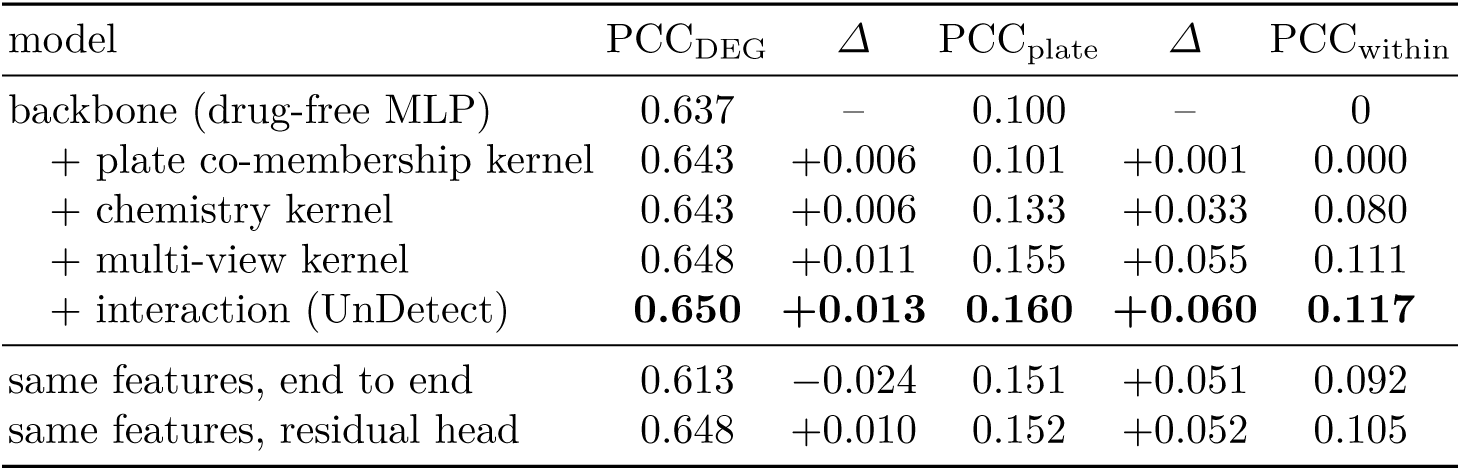
Ablation on the drug-blind benchmark, mean of five folds, all rows on one backbone run. *Δ*: mean paired difference to the backbone; every difference has the same sign in 5/5 folds with bootstrap intervals over test compounds excluding zero, except the plate co-membership kernel on PCC_plate_ (above zero in 2/5). Same features: UnDetect’s compound features given to the backbone network next to the control, or to a two-layer network on the backbone’s residuals.

| model | PCC<sub>DEG</sub> | *Δ* | PCC<sub>plate</sub> | *Δ* | PCC<sub>within</sub> |
|---|---|---|---|---|---|
| backbone (drug-free MLP) | 0.637 | – | 0.100 | – | 0 |
| + plate co-membership kernel | 0.643 | +0.006 | 0.101 | +0.001 | 0.000 |
| + chemistry kernel | 0.643 | +0.006 | 0.133 | +0.033 | 0.080 |
| + multi-view kernel | 0.648 | +0.011 | 0.155 | +0.055 | 0.111 |
| + interaction (UnDetect) | **0.650** | **+0.013** | **0.160** | **+0.060** | **0.117** |
| same features, end to end | 0.613 | −0.024 | 0.151 | +0.051 | 0.092 |
| same features, residual head | 0.648 | +0.010 | 0.152 | +0.052 | 0.105 |

**Table 5:** PCC_within_ by the compound information beyond structure (drug-blind benchmark, five folds pooled; compounds on plates with at least two test compounds; the backbone scores 0).

| information | compounds | DeepCE | TranSiGen | XPert | UnDetect | UnDetect + XPert |
|---|---|---|---|---|---|---|
| structure only | 4,615 | 0.017 | 0.061 | 0.068 | 0.071 | **0.083** |
| + targets, mechanism | 1,575 | 0.035 | 0.091 | 0.103 | 0.112 | **0.128** |
| + Cell Painting | 573 | 0.030 | 0.074 | 0.079 | **0.159** | 0.152 |
| + both | 1,353 | 0.034 | 0.095 | 0.120 | 0.170 | **0.175** |
| all | 8,116 | 0.027 | 0.079 | 0.092 | 0.117 | **0.126** |

Fitting the compound after the context is what keeps both. Given to the benchmark network next to the control and trained end to end, UnDetect’s compound features lower PCC_DEG_ below ignoring the compound; fitted to the backbone’s residuals by a neural head, they keep the compound signal and recover the context, and UnDetect remains ahead on all three scores.

Every compound view carries response information, and chemistry and Cell Painting each carry information that no other view supplies (fig. 3a). Each view alone gives about half of UnDetect’s gain in PCC_plate_, and the four together beat every single view in all folds. Removing or permuting chemistry or Cell Painting lowers the score in every fold, whereas targets and mechanism are largely redundant with each other. Supplementary Table 14 gives every view ablation.

**Fig. 3:**
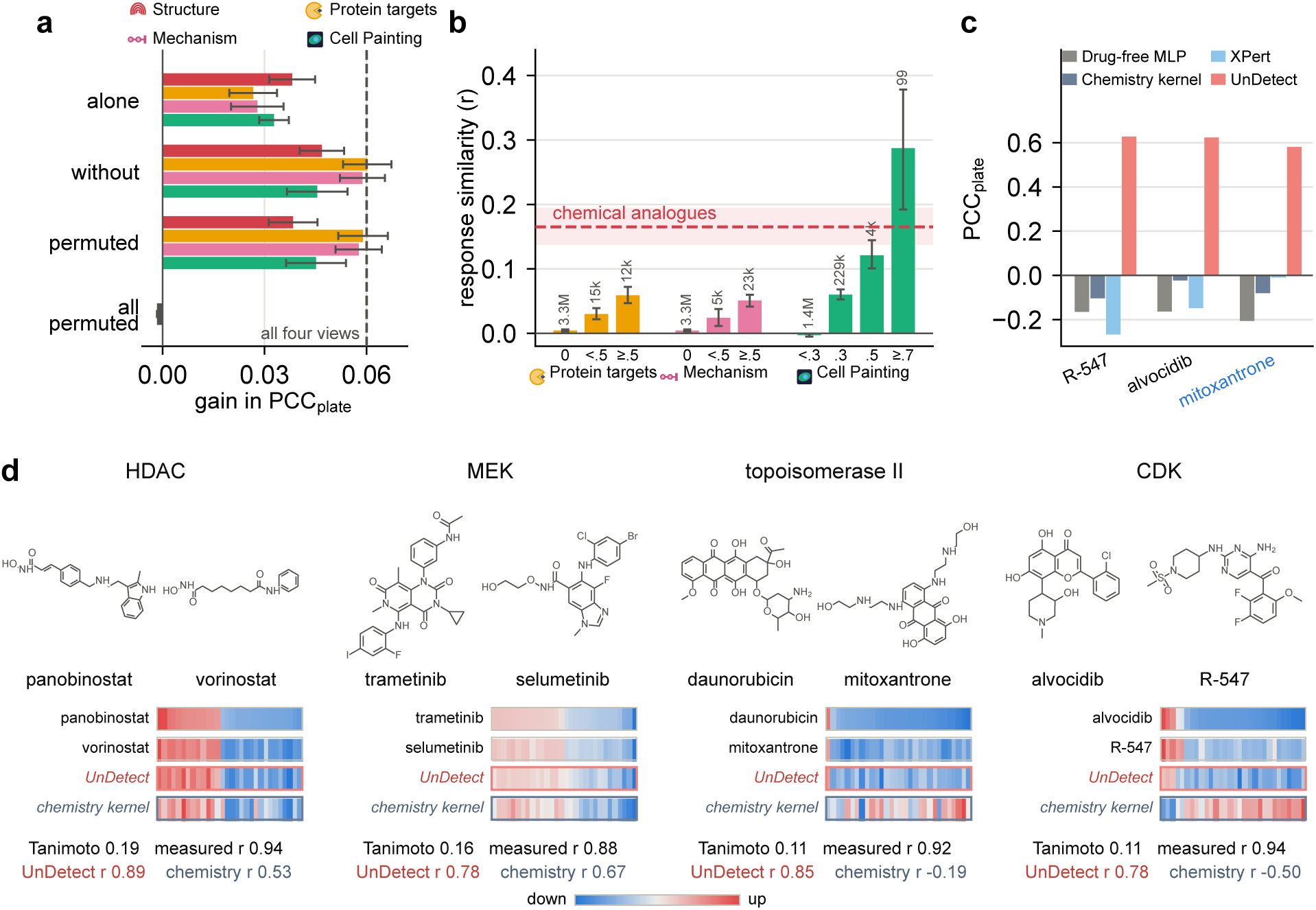
What the compound views add beyond chemistry (drug-blind benchmark). (a) Every view carries response information: gain in PCC_plate_ over the drug-free backbone with one view alone, with all views but one, with one view’s compound labels permuted, or with all four permuted; the dashed line marks all four views. (b) Chemically unlike compounds (Tanimoto *<* 0.25) respond alike when their targets, mechanisms or, most of all, Cell Painting profiles are similar; the band marks chemical analogues (Tanimoto *≥* 0.6). (c) Where structure points to the wrong neighbors, the views recover the response: the three test compounds on which UnDetect gains most over XPert and the chemistry kernel, among the 337 test compounds with common names, each scored while held out. (d) Four chemically unlike pairs that respond alike, one per mechanism, with the signatures of both and the second compound’s signature as predicted while held out; red up, blue down. Error bars show the s.d. over folds or permutations in (a) and 95% intervals over compounds in (b). settings, with intervals above zero, whereas on Tahoe-100M, with few compounds per line, validation mostly switches it off. On DeepCE’s benchmark a neural network on chemistry alone is the better regressor (0.511 against 0.491), so UnDetect’s lead comes from combining partially observed views; across ten compounddisjoint resplits, UnDetect beats chemistry-only kernel regression in every fold, and its cross-line coupling and its normalization each help in nine (Supplementary Table 10).

UnDetect’s gain grows with what is known about a compound (table 5). With structure alone, UnDetect is level with XPert, whose compound embedding also draws on drug–target links, on PCC_within_ and above TranSiGen and DeepCE, and on PCC_DEG_ it is above all five published models with released predictions. Its lead grows with targets and mechanism, and with Cell Painting it reaches up to twice XPert’s score. The ensemble of the two helps most where compound information is scarce. Supplementary Table 16 gives PCC_plate_ by view coverage.

On LINCS P2-24 the target kernel beats a kernel over permuted target annotations in all 7 lines, whereas a kernel over the targets’ one-hop STRING [31] neighborhoods, the gene-network prior common in perturbation models [24,27,9,12], is no better than the same kernel on a degree-preserving rewired [20] network. UnDetect therefore uses pharmacological and phenotypic views and no interaction network; Supplementary Table 20 gives the null tests.

On LINCS and Tahoe-100M the shared term alone is below the per-line kernel in every setting, so both parts are needed (table 3); the interaction with the cell line as context adds 0.0005 to 0.0014 in all four LINCS

### 3.5 Compound case study

Pairs of compounds show why the views help (fig. 3b, d). Among chemically unlike pairs, response similarity rises with every other view, and pairs with similar Cell Painting profiles (*≥* 0.7) respond more alike than chemical analogues do (Pearson *r* 0.29, 95% interval 0.19–0.38, against 0.17, 0.14–0.20), and still do when they share no target or mechanism (0.26); panobinostat and vorinostat, for instance, share little structure yet respond almost identically. The query of fig. 1 shows the mechanism at work. By structure alone mitoxantrone is closest to vorinostat; the fused kernel pairs it with daunorubicin, which shares its target and mechanism. Held out from training, it is predicted by UnDetect with PCC_plate_ 0.58, whereas the chemistry kernel and XPert recover none of its compound-specific response. UnDetect improves on the chemistry kernel for 72% of the named test compounds, and its largest gains come where structure points to the wrong neighbors, as for R-547, alvocidib and mitoxantrone (fig. 3c). For the second compound of each pair in fig. 3d, UnDetect’s prediction while held out matches the measured signature closely, unlike that of the chemistry kernel. Supplementary Table 13 lists further chemically unlike pairs that respond alike.

## 4 Discussion

Current models for unprofiled compounds score well on the drug-blind benchmark for a reason unrelated to the compound: the paired control identifies the plate, and the plate predicts most of the response. Fitting the compound after this context, UnDetect attains, without any measurement of the compound, 60% of the gain of a reference that uses the compound’s measured responses in other cell lines, and its average with XPert 86%. As for genetic perturbations [1,35], a simple model with the right decomposition, fitted in minutes, is a strong baseline for end-to-end models.

The compound information that generalizes is pharmacological and phenotypic (table 5), which fits two uses. In drug repurposing, approved and investigational drugs already come with annotated targets and mechanisms. For new chemistry, one Cell Painting assay supplies the most informative view at a fraction of the effort of transcriptional profiling across cell lines, a practical first experiment; profiles from a single cell line (A549) already generalize to 164 lines. Protein-interaction priors, in contrast, add nothing beyond node degree, so network priors should be calibrated against rewired nulls.

Drug-blind evaluations should report a plate-aware score, such as PCC_within_, or keep plate-mates together in the split; otherwise a model that ignores the compound can rank first. All cell lines in our evaluations are seen in training, and one assay (A549 Cell Painting) supplies the phenotypic view; for unseen cell lines [30,21], the interaction through the control profile is a natural starting point.

## Author contributions

Y.L. designed and implemented the method, built the benchmarks, made the figures and wrote the paper. H.L. made figures, reviewed the paper and contributed to the method discussions.

K.P. performed the downstream analyses and the analysis of the results. J.W. conceived and supervised the project, set its direction and guided the discussions.

## Appendix

### **A.1** Implementation details

*Backbone.* On the drug-blind benchmark the backbone is the drug-free network of Bai et al. [3]: three

hidden layers of 2,048 ReLU units with dropout 0.1, loss 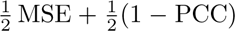, Adam with learning rate 10*^−^*^4^, batch 512 and gradient norm clipped at 1, at most 500 epochs, early stopping on the PCC_DEG_ of validation compounds with patience 50. It is trained on training compounds only and then frozen. On the Level-5 (LINCS, DeepCE) and Tahoe-100M benchmarks the control has already been removed, and the only compound-free inputs are the cell line and, on DeepCE’s benchmark, the dose; the least-squares backbone is then the training mean of each line (and dose), and all compound information enters through the kernel terms. Retrained under two further seeds, in an earlier configuration (kernel terms with a neural head, no interaction), the five-fold mean PCC_DEG_ of the backbone and of the model on it changed by at most 0.0003, and the gain over the drug-free network was positive in all 15 fold–seed runs.

*Kernel normalization* Structure receives a view weight of 0.28 to 0.31 in every fold, so every diagonal entry of *Δ* is positive; the implementation clamps it at 10*^−^*^12^, so a compound whose views all had zero weight would keep a zero row and be predicted by the backbone.

*Selection grids* are listed in table 6. On the drug-blind benchmark *λ* is chosen with the shared term alone, then (*a, b*) jointly, then (*κ, q*), each on validation compounds. On LINCS and Tahoe-100M, *λ* is chosen per fold (per cell line for the line-specific term) and (*a, b*) per cell line and fold, by compound-grouped inner cross-validation.

**Table 6:** Hyperparameter grids. DeepCE’s benchmark fits one joint kernel ridge regression without the interaction; *σ* = 0 and *σ* = *∞* are the limits of the dose kernel (a separate fit per dose, and similarity independent of dose).

| benchmark | parameter grid |  |
|---|---|---|
| drug-blind | *λ* | 0.1, 0.3, 1, 3, 10, 30, 100 |
|  | *a*, *b* | 0, 0.25, 0.5, 0.75, 1, 1.25, 1.5, 2 |
|  | *κ* | 10<sup>−3</sup>, 10<sup>−2</sup>, 10<sup>−1</sup>, 1, 10, ∞ |
|  | *q* | 5, 10, 20 |
| LINCS, Tahoe-100M | *λ* | 10<sup>−2</sup>, 10<sup>−1</sup>, 1, 10, 10<sup>2</sup>, 10<sup>3</sup> |
|  | *a* | 0, 0.25, 0.5, 0.75, 1, 1.5, 2 |
|  | *b* | 0, 0.25, 0.5, 0.75, 1 |
|  | *κ* | 10<sup>−3</sup>, 10<sup>−2</sup>, 10<sup>−1</sup>, 1, 10, ∞ |
| DeepCE | *λ* | 0.01, 0.03, 0.1, 0.3, 1, 3, 10, 30 |
|  | *ρ* | 0, 0.1, 0.25, 0.5, 1, 2, 4 |
|  | *σ* | 0, 0.25, 0.5, 1, 2, 4, ∞ |

### **A.2** Drug-blind benchmark, per fold

**Table 7:**
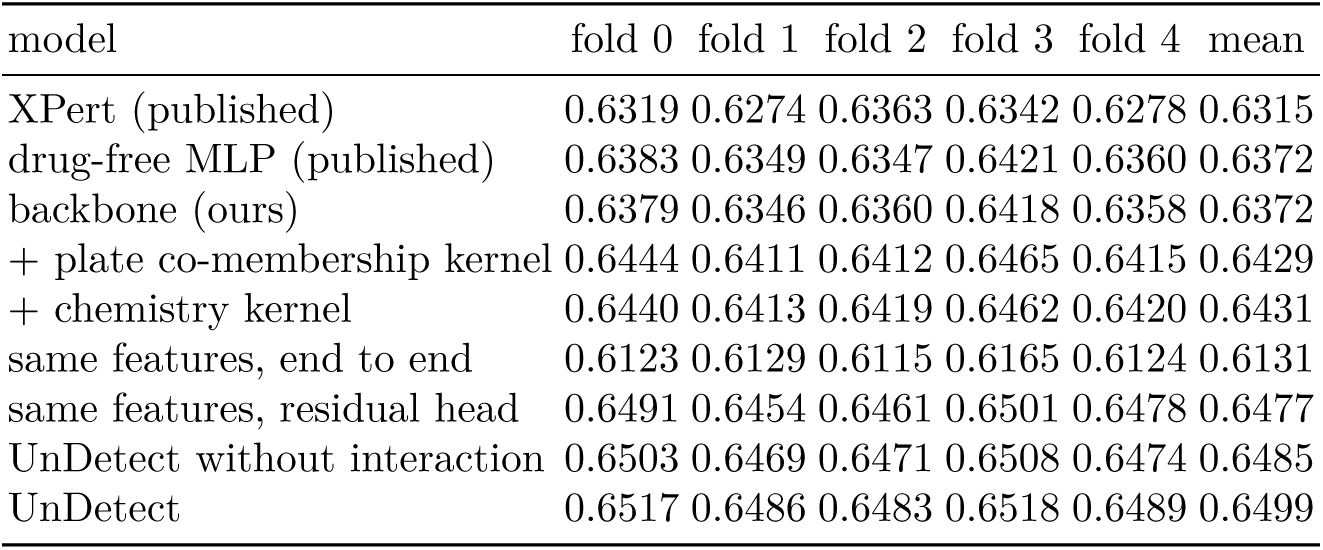
Drug-blind benchmark, PCC_DEG_ per fold (test compounds). Published rows from the predictions released by Bai et al. [3]; all other rows are our runs on one backbone (seed 1729).

| model | fold 0 | fold 1 | fold 2 | fold 3 | fold 4 | mean |
|---|---|---|---|---|---|---|
| XPert (published) | 0.6319 | 0.6274 | 0.6363 | 0.6342 | 0.6278 | 0.6315 |
| drug-free MLP (published) | 0.6383 | 0.6349 | 0.6347 | 0.6421 | 0.6360 | 0.6372 |
| backbone (ours) | 0.6379 | 0.6346 | 0.6360 | 0.6418 | 0.6358 | 0.6372 |
| + plate co-membership kernel | 0.6444 | 0.6411 | 0.6412 | 0.6465 | 0.6415 | 0.6429 |
| + chemistry kernel | 0.6440 | 0.6413 | 0.6419 | 0.6462 | 0.6420 | 0.6431 |
| same features, end to end | 0.6123 | 0.6129 | 0.6115 | 0.6165 | 0.6124 | 0.6131 |
| same features, residual head | 0.6491 | 0.6454 | 0.6461 | 0.6501 | 0.6478 | 0.6477 |
| UnDetect without interaction | 0.6503 | 0.6469 | 0.6471 | 0.6508 | 0.6474 | 0.6485 |
| UnDetect | 0.6517 | 0.6486 | 0.6483 | 0.6518 | 0.6489 | 0.6499 |

**Table 8:**
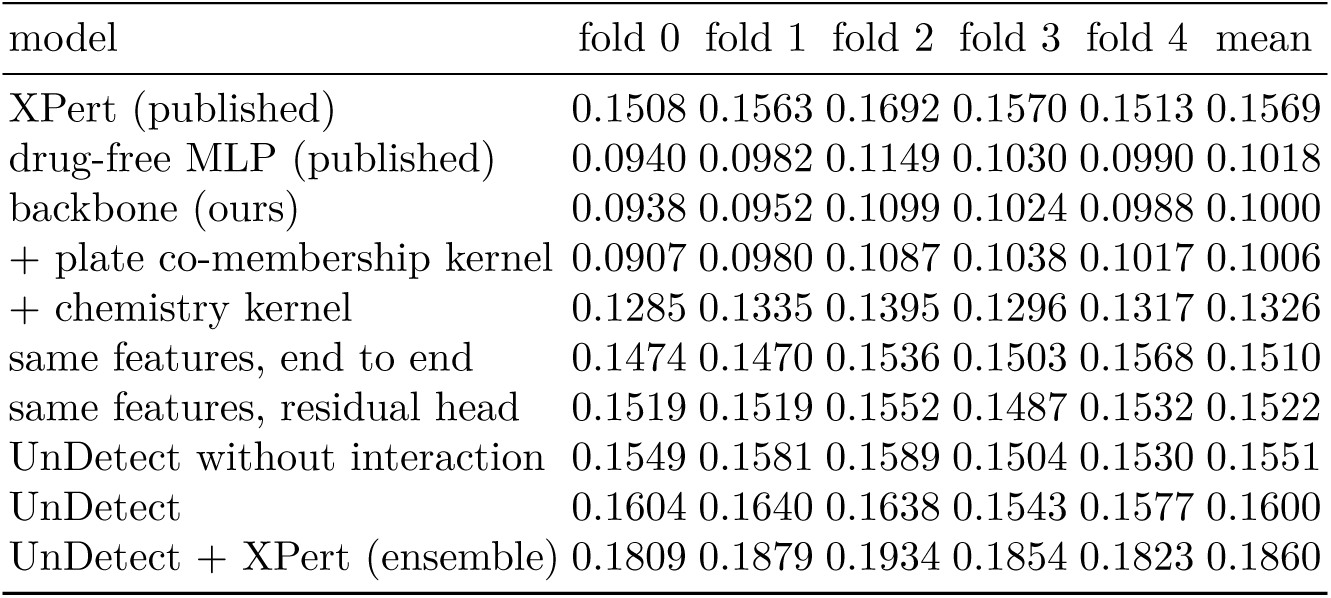
Drug-blind benchmark, PCC_plate_ per fold (test samples with a training plate-mate).

| model | fold 0 | fold 1 | fold 2 | fold 3 | fold 4 | mean |
|---|---|---|---|---|---|---|
| XPert (published) | 0.1508 | 0.1563 | 0.1692 | 0.1570 | 0.1513 | 0.1569 |
| drug-free MLP (published) | 0.0940 | 0.0982 | 0.1149 | 0.1030 | 0.0990 | 0.1018 |
| backbone (ours) | 0.0938 | 0.0952 | 0.1099 | 0.1024 | 0.0988 | 0.1000 |
| + plate co-membership kernel | 0.0907 | 0.0980 | 0.1087 | 0.1038 | 0.1017 | 0.1006 |
| + chemistry kernel | 0.1285 | 0.1335 | 0.1395 | 0.1296 | 0.1317 | 0.1326 |
| same features, end to end | 0.1474 | 0.1470 | 0.1536 | 0.1503 | 0.1568 | 0.1510 |
| same features, residual head | 0.1519 | 0.1519 | 0.1552 | 0.1487 | 0.1532 | 0.1522 |
| UnDetect without interaction | 0.1549 | 0.1581 | 0.1589 | 0.1504 | 0.1530 | 0.1551 |
| UnDetect | 0.1604 | 0.1640 | 0.1638 | 0.1543 | 0.1577 | 0.1600 |
| UnDetect + XPert (ensemble) | 0.1809 | 0.1879 | 0.1934 | 0.1854 | 0.1823 | 0.1860 |

### **A.3** DeepCE benchmark

*Reproduction.* We ran DeepCE from its released code (commit 9b0d04f) on the published split (1,895, 544 and 496 training, validation and test signatures; 92 test compounds) with its STRING gene features, selecting the epoch on the validation set. Three seeds give 0.481, 0.456 and 0.428 per-signature Pearson, mean 0.455, against 0.491 reported by Pham et al. [24]. In fig. 2c and table 9, “DeepCE” is this reproduction; the *k*NN (0.390), neural network (0.393) and DeepCE without attention (0.442) are the values reported in that paper. The headline comparison is with the reported 0.491, which favors DeepCE. Kernel regression on chemistry alone scores 0.4911, within 0.001 of the reported DeepCE value of 0.4907; the two are separate measurements. The compound views of the 92 test compounds are chemistry for all, targets and MOA for 62 and Cell Painting for 72.

**Table 9:** DeepCE benchmark, per-signature Pearson on the published test split. UnDetect: kernel terms, without the interaction. MLP: two hidden layers on the same inputs and target, tuned on the validation set, mean of thr<u>ee seeds.</u>

| Baselines |  | Kernel regression |  |
|---|---|---|---|
| model | Pearson | model | Pearson |
| mean per line and dose | 0.389 | KRR, chemistry | 0.491 |
| Morgan *k*NN, tuned | 0.451 | UnDetect, chem. + targets + MOA | 0.567 |
| DeepCE (reproduced) | 0.455 | UnDetect, all views | **0.602** |
| MLP, chemistry | 0.511 |  |  |
| MLP, all views | 0.544 |  |  |

*Resplits.* All 2,935 signatures were pooled and split five ways by compound, twice with different seeds; each fold selects its hyperparameters on one fifth of its training compounds with the grids of table 6 (table 10).

**Table 10:**
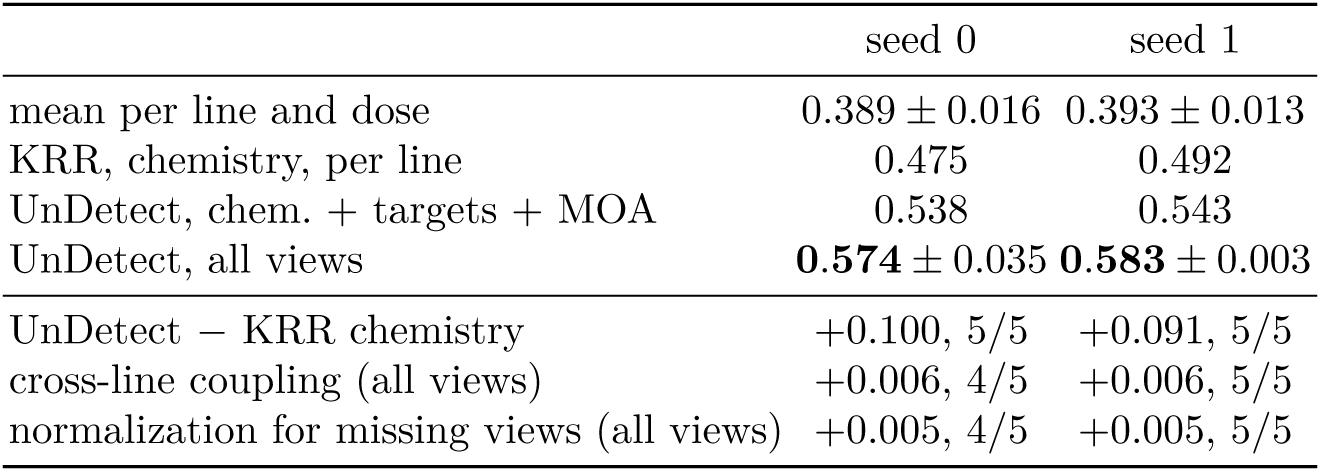
DeepCE data, five compound-disjoint folds for each of two seeds: per-signature Pearson (mean *±* s.d. over folds) and fold-paired differences (mean; folds where the difference is positive).

### **A.4** Interaction variants

**Table 11:**
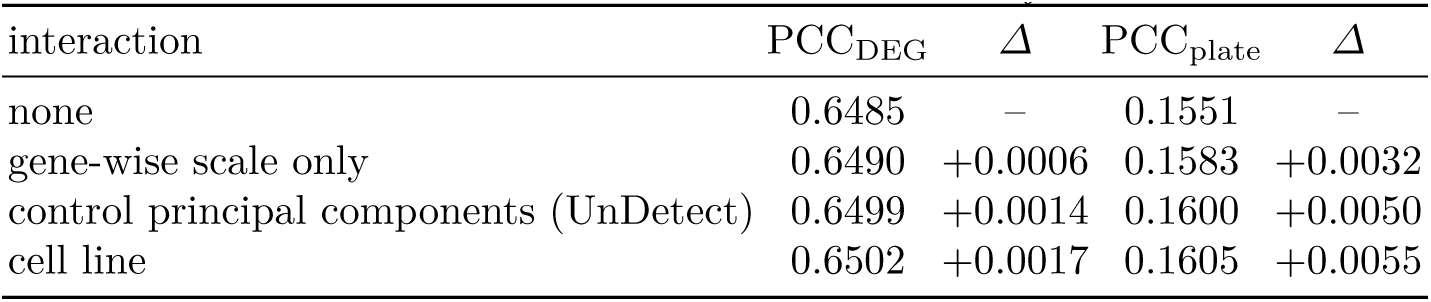
Interaction variants on the drug-blind benchmark, five-fold means. *Δ*: paired difference to UnDetect without interaction; every *Δ* is positive in 5/5 folds with bootstrap intervals above zero. Gene-wise scale: *γ_j_* = 0 in eq. (1). Cell line: one scale per cell line and gene instead of 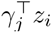.

| interaction | PCC<sub>DEG</sub> | *Δ* | PCC<sub>plate</sub> | *Δ* |
|---|---|---|---|---|
| none | 0.6485 | – | 0.1551 | – |
| gene-wise scale only | 0.6490 | +0.0006 | 0.1583 | +0.0032 |
| control principal components (UnDetect) | 0.6499 | +0.0014 | 0.1600 | +0.0050 |
| cell line | 0.6502 | +0.0017 | 0.1605 | +0.0055 |

The interaction strength chosen on validation compounds is *κ* = 1 in every fold, with *q* = 20 principal components in four folds and *q* = 10 in one.

### **A.5** Published drug-input ablation

**Table 12:** The published models with their drug input and with it replaced by zeros, fold means from the predictions released by Bai et al. [3]. PRnet and PertDiT are released in their own normalized expression space, so no PCC_plate_ is computed for them; CIGER’s zeroed predictions are constant within a plate. PertDiT’s difference is *−*0.01<u>248, quoted as 0.012 in the main text.</u>

| model | PCC<sub>DEG</sub> |  |  | PCC<sub>plate</sub> |  |  |
|---|---|---|---|---|---|---|
|  | with drug | zeroed | difference | with drug | zeroed | difference |
| CIGER [23] | 0.2456 | 0.2485 | −0.0028 | 0.0108 | – | – |
| MultiDCP [36] | 0.2545 | 0.2492 | +0.0052 | 0.0244 | 0.0230 | +0.0014 |
| PRnet [25] | 0.2771 | 0.2764 | +0.0007 | – | – | – |
| PertDiT [17] | 0.3701 | 0.3825 | −0.0125 | – | – | – |
| DeepCE [24] | 0.3947 | 0.3834 | +0.0113 | 0.0553 | 0.0474 | +0.0079 |
| TranSiGen [32] | 0.6144 | 0.6057 | +0.0088 | 0.1440 | 0.1012 | +0.0428 |
| XPert [12] | 0.6315 | 0.6313 | +0.0002 | 0.1569 | 0.1064 | +0.0505 |

### **A.6** Chemically unlike compounds that respond alike

**Table 13:** Pairs with Morgan Tanimoto below 0.25 in the top 0.5% of the fused kernel, ranked by response similarity (Pearson correlation of plate-centered signatures), one or two per mechanism. Fused: normalized multi-view kernel. An aste<u>risk marks a compound without target or mechanism ann</u>otation in the benchmark.

| compound | compound | Tanimoto | fused | response *r* |
|---|---|---|---|---|
| panobinostat | vorinostat | 0.19 | 0.50 | 0.94 |
| alvocidib | R-547<sup>*</sup> | 0.11 | 0.25 | 0.94 |
| NVP-BEZ235 | PI-103 | 0.13 | 0.54 | 0.92 |
| daunorubicin | mitoxantrone | 0.11 | 0.43 | 0.92 |
| AZD-8055 | PI-103 | 0.20 | 0.25 | 0.91 |
| anguidine | emetine | 0.09 | 0.33 | 0.91 |
| MG-132 | ixazomib | 0.21 | 0.61 | 0.90 |
| orciprenaline | procaterol | 0.20 | 0.70 | 0.90 |
| trametinib | selumetinib | 0.16 | 0.45 | 0.88 |
| triptolide<sup>*</sup> | alvocidib | 0.10 | 0.26 | 0.87 |
| BVD-523<sup>*</sup> | TAK-733 | 0.14 | 0.31 | 0.86 |

### **A.7** View ablation

**Table 14:** View ablation on the drug-blind benchmark: UnDetect with the interaction, on the backbone of table 4, mean of five folds. Permuted: the view’s compound labels shuffled among the compounds that have it. *Δ*: PCC_plate_ difference to all four views; folds: number of folds with the bootstrap interval over test compounds below zero.

| views | PCC<sub>DEG</sub> | PCC<sub>plate</sub> | *Δ* | folds |
|---|---|---|---|---|
| none (backbone) | 0.6372 | 0.1000 | −0.0600 | 5 |
| all four | 0.6499 | 0.1600 | – | – |
| chemistry alone | 0.6441 | 0.1381 | −0.0219 | 5 |
| targets alone | 0.6437 | 0.1267 | −0.0334 | 5 |
| mechanism alone | 0.6438 | 0.1279 | −0.0322 | 5 |
| Cell Painting alone | 0.6430 | 0.1328 | −0.0273 | 5 |
| without chemistry | 0.6475 | 0.1469 | −0.0132 | 5 |
| without targets | 0.6497 | 0.1602 | +0.0002 | 0 |
| without mechanism | 0.6494 | 0.1588 | −0.0012 | 1 |
| without Cell Painting | 0.6470 | 0.1455 | −0.0145 | 5 |
| chemistry permuted | 0.6460 | 0.1384 | −0.0216 | 5 |
| targets permuted | 0.6492 | 0.1590 | −0.0011 | 2 |
| mechanism permuted | 0.6489 | 0.1577 | −0.0023 | 4 |
| Cell Painting permuted | 0.6470 | 0.1452 | −0.0149 | 5 |

### **A.8** Controls for the compound terms

Permutation null: one permutation of compound identities, drawn within each group of compounds that have the same views, is applied to all four view kernels at once; every sample of a compound therefore receives the same permuted compound, and plates, cell lines, samples and view coverage are unchanged. The view weights, the kernel terms with their *λ* and scales, and the interaction (*κ*, *q*) are refitted; the backbone is fixed. Five permutations. Compound-stage cross-fitting: on the fixed backbone, the training-sample kernel terms that feed the interaction are predicted for each fifth of the training compounds from the other four fifths, with the view weights and fused kernel re-estimated each time; *λ* and the scales are those of the main model.

**Table 15:** Controls on the drug-blind benchmark, per fold. Permuted: UnDetect PCC_plate_ with jointly permuted compound views, mean of five permutations (backbone in parentheses). Interaction gain: UnDetect minus its kernel terms, PCC_plate_, with leave-one-compound-out training terms (main model) and with compound-stage cross-fitting; 95% bootstrap intervals over test compounds.

| fold | UnDetect | permuted (backbone) | interaction, leave-one-compound-out | interaction, cross-fitted |
|---|---|---|---|---|
| 0 | 0.160 | 0.092 (0.094) | +0.0055 [0.0036, 0.0076] | +0.0055 [0.0036, 0.0076] |
| 1 | 0.164 | 0.094 (0.095) | +0.0059 [0.0042, 0.0078] | +0.0060 [0.0042, 0.0077] |
| 2 | 0.164 | 0.109 (0.110) | +0.0049 [0.0039, 0.0060] | +0.0050 [0.0039, 0.0061] |
| 3 | 0.154 | 0.100 (0.102) | +0.0039 [0.0019, 0.0057] | +0.0039 [0.0019, 0.0058] |
| 4 | 0.158 | 0.097 (0.099) | +0.0047 [0.0033, 0.0061] | +0.0047 [0.0033, 0.0061] |
| mean | 0.160 | 0.098 (0.100) | +0.0050 | +0.0050 |

### A.9 Sharing across cell lines by coverage and gain by compound information

**Fig. 4:**
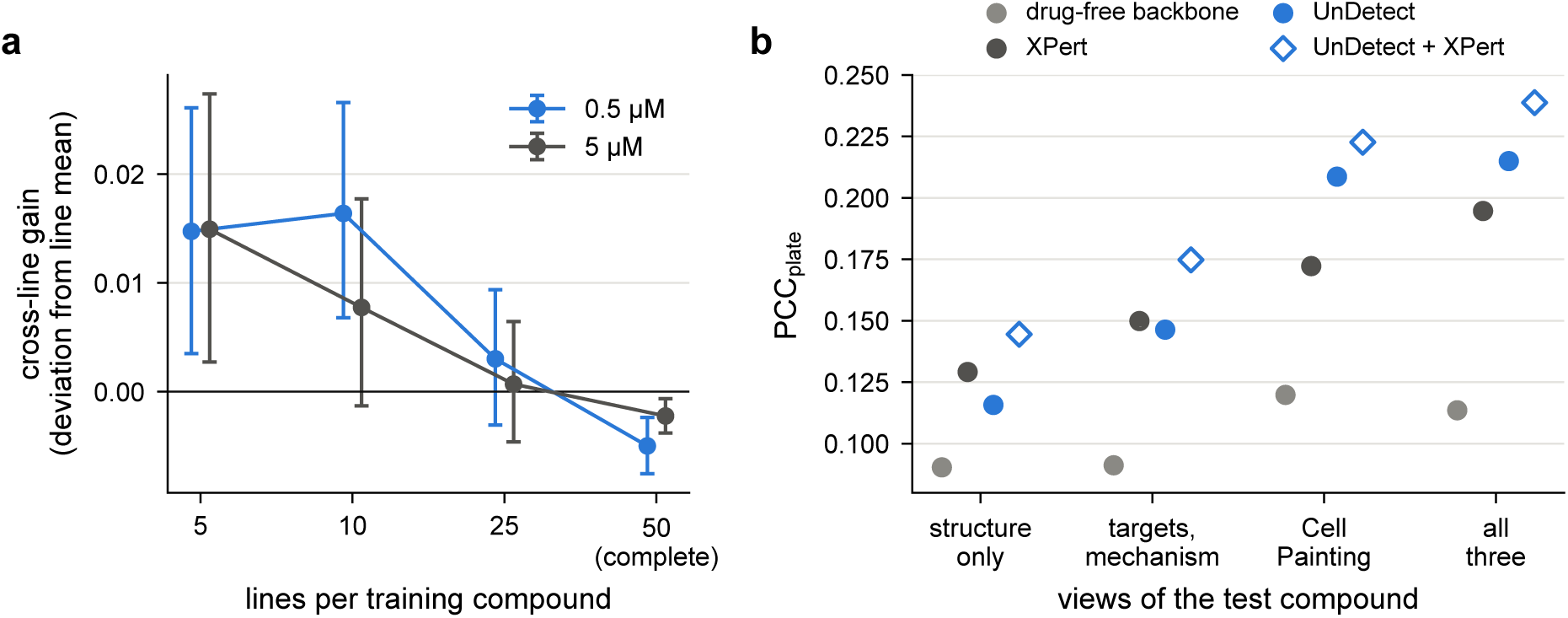
(a) Tahoe-100M: gain of the cross-line term over the per-line kernel on the correlation of deviations from the line mean, by the number of lines in which each training compound is kept (50: the complete grid); test compounds keep all lines; bars: 95% bootstrap intervals over compound groups. (b) Drug-blind benchmark: PCC_plate_ by the views a test compound has (numbers in table 16).

**Table 16:** Drug-blind benchmark by the views a test compound has (targets and mechanism have identical coverage), PCC_plate_, five folds pooled, compounds with a training plate-mate. Ensemble: UnDetect + XPert as in table 2<u>. Differences with 95% bootstrap intervals over test compounds.</u>

| views | compounds | backbone | UnDetect | XPert | UnDetect − backbone | ensemble − XPert |
|---|---|---|---|---|---|---|
| structure only | 4,693 | 0.090 | 0.116 | 0.129 | +0.025 [0.021, 0.030] | +0.015 [0.014, 0.017] |
| targets, mechanism | 1,591 | 0.091 | 0.146 | 0.150 | +0.055 [0.041, 0.072] | +0.025 [0.020, 0.030] |
| Cell Painting | 578 | 0.120 | 0.209 | 0.172 | +0.089 [0.073, 0.105] | +0.050 [0.041, 0.062] |
| all three | 1,355 | 0.114 | 0.215 | 0.195 | +0.101 [0.085, 0.118] | +0.044 [0.037, 0.052] |

### **A.10** PCC_plate_ by the number of training plate-mates

On PCC_within_, which uses no plate-mean estimate, UnDetect leads XPert in every bin, by 0.046 with at most two plate-mates and by 0.010 with more than fifty, all intervals above zero; XPert’s lead on PCC_plate_ for plates with few plate-mates does not carry over to it.

**Fig. 5:**
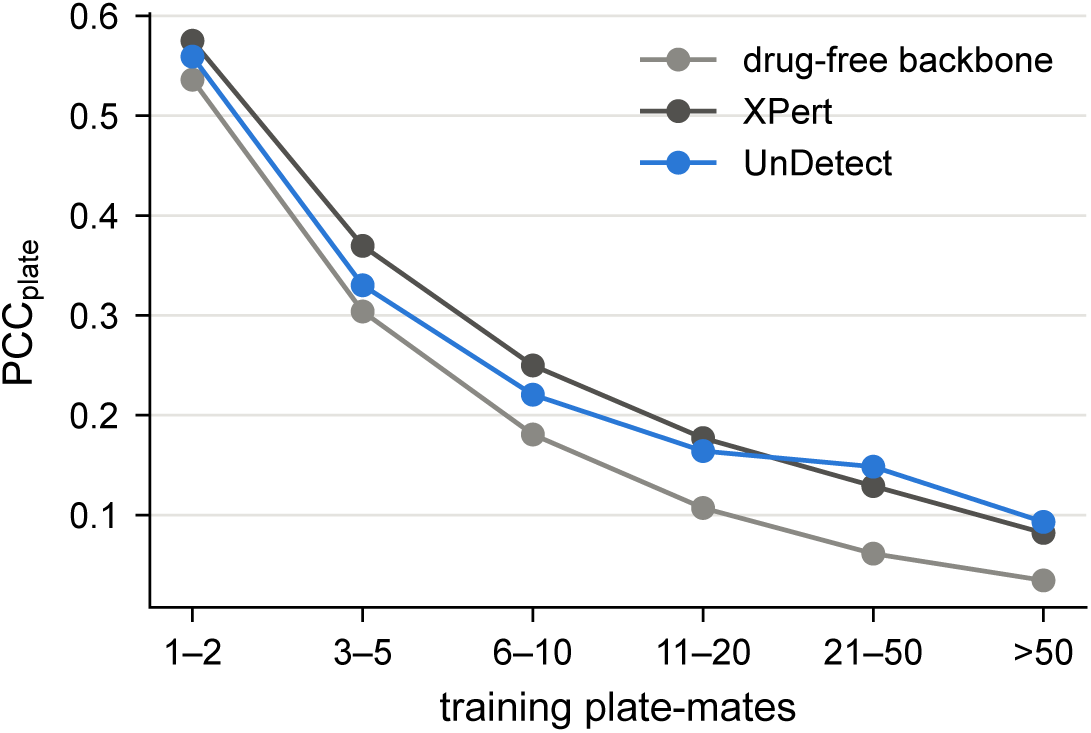
Drug-blind benchmark, PCC_plate_ by the number of training samples that share the test sample’s control (numbers and intervals in table 17).

**Table 17:** Drug-blind benchmark, PCC_plate_ by the number of training samples that share the test sample’s contr<u>ol (plate-mates), fi</u>v<u>e folds pooled. Differenc</u>e<u>s with 95% bootstrap intervals over test compounds</u>.

| plate-mates | samples | backbone | UnDetect | XPert | UnDetect − backbone | UnDetect − XPert |
|---|---|---|---|---|---|---|
| 1–2 | 3,731 | 0.536 | 0.559 | 0.575 | +0.023 [0.019, 0.028] | −0.016 [−0.021, −0.011] |
| 3–5 | 3,650 | 0.304 | 0.330 | 0.370 | +0.026 [0.022, 0.031] | −0.040 [−0.045, −0.034] |
| 6–10 | 4,988 | 0.181 | 0.221 | 0.250 | +0.040 [0.035, 0.045] | −0.029 [−0.035, −0.024] |
| 11–20 | 8,558 | 0.107 | 0.164 | 0.177 | +0.057 [0.052, 0.062] | −0.013 [−0.018, −0.007] |
| 21–50 | 16,298 | 0.061 | 0.148 | 0.129 | +0.087 [0.075, 0.100] | +0.019 [0.010, 0.029] |
| more than 50 | 33,740 | 0.035 | 0.093 | 0.082 | +0.059 [0.051, 0.068] | +0.011 [0.005, 0.017] |

### **A.11** Within-plate score

For each fold, test samples are grouped by their control row, which identifies the plate, and averaged per (plate, compound) when a compound occurs more than once on a plate (at most 10 rows per fold). Plates with at least two test compounds are kept: 1,525 to 1,575 per fold, covering 86% of test samples, including samples without a training plate-mate. For unit (*p, d*),

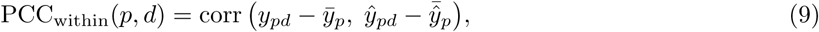

where *y̅_p_* and *y̅*^*_p_* are the means of truth and prediction over the plate’s test compounds; the score is averaged over units. A prediction that is constant within a plate has a zero centered vector, for which the correlation is undefined; by convention it scores 0, the value of a prediction without compound information; the backbone’s predictions are exactly constant, and centered predictions below 10*^−^*^3^ in absolute value (the published MLP, stored at half precision) are set to zero. Intervals: 2,000 bootstrap replicates over test compounds per fold (table 18). Compounds on one plate share its centering, so we also resampled whole plates: UnDetect, its interaction gain and its lead over XPert keep intervals above zero in every fold. Resampling plates and compounds together, the lead over XPert is above zero in four of five folds (lowest bound *−*0.0004).

**Table 18:** PCC_within_ on the drug-blind benchmark, per fold, and paired differences (mean of five folds; folds with the 95% interval above zero).

| model | fold 0 | fold 1 | fold 2 | fold 3 | fold 4 | mean |
|---|---|---|---|---|---|---|
| backbone, published MLP | 0 | 0 | 0 | 0 | 0 | 0 |
| XPert | 0.087 | 0.098 | 0.102 | 0.080 | 0.094 | 0.092 |
| XPert, drug input zeroed | 0 | 0 | 0 | 0 | 0 | 0 |
| TranSiGen | 0.075 | 0.086 | 0.083 | 0.074 | 0.076 | 0.079 |
| UnDetect | 0.120 | 0.123 | 0.110 | 0.111 | 0.119 | **0.117** |
| UnDetect + XPert | 0.124 | 0.133 | 0.126 | 0.118 | 0.130 | 0.126 |
| UnDetect − XPert | intervals above zero in 5/5 folds |  |  |  |  | +0.025 |
| ensemble − UnDetect | intervals above zero in 4/5 folds |  |  |  |  | +0.009 |
| UnDetect − same features, residual head | intervals above zero in 5/5 folds |  |  |  |  | +0.012 |
| interaction (UnDetect − without it) | intervals above zero in 5/5 folds |  |  |  |  | +0.006 |

### **A.12** Within-plate self-retrieval

Units and centering as for PCC_within_ (appendix A.11): 67,680 (plate, compound) units of 8,116 test compounds on plates with at least two test compounds, five folds pooled. For each unit, the measured signatures of the plate’s *n* test compounds are ranked by their correlation with the compound’s predicted signature (ties averaged). Top-1: the compound’s own signature ranks first (chance: the mean of 1*/n*); normalized rank: (rank *−* 1)*/*(*n −* 1), 0 best and 0.5 by chance; predictions constant within a plate (backbone, XPert with its drug input zeroed) score exactly 0.5. Differences: 95% bootstrap intervals over test compounds (table 19).

**Table 19:**
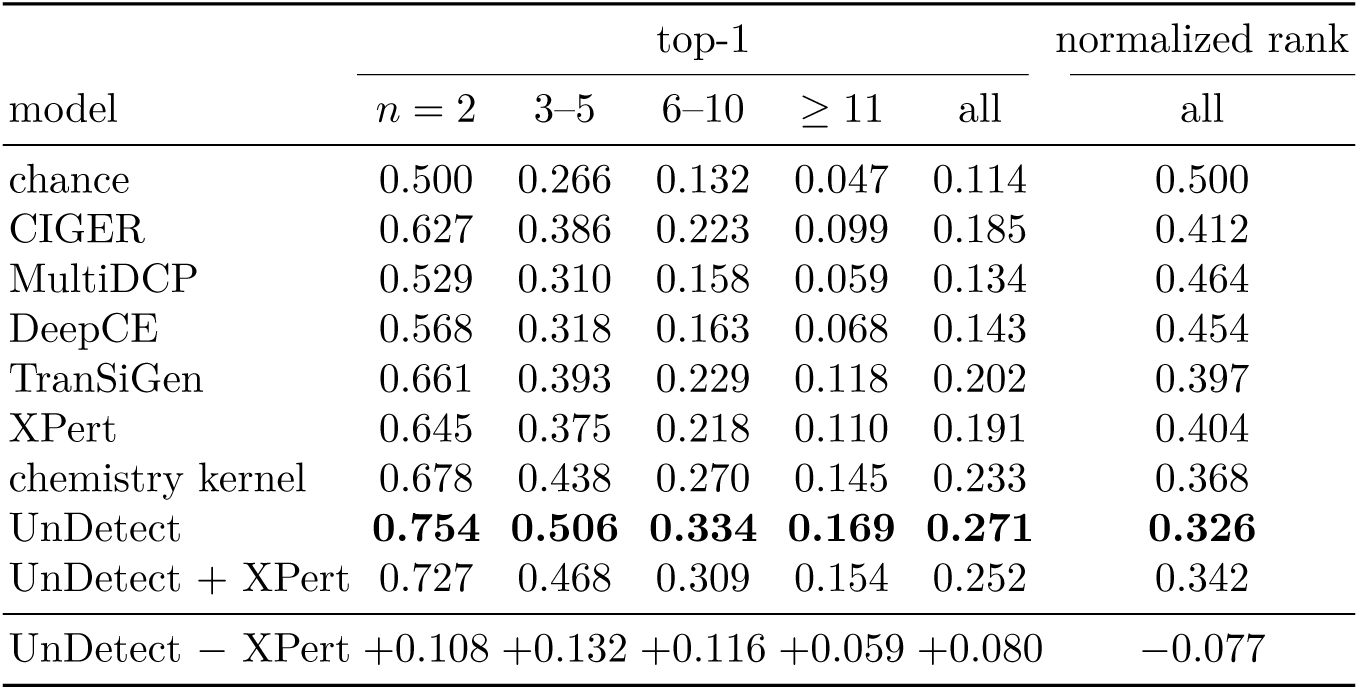
Within-plate self-retrieval on the drug-blind benchmark, by plate size *n* (number of test compounds on the plate). The 95% intervals of UnDetect *−* XPert exclude zero in every column (all: top-1 [0.074, 0.087], normalized rank [*−*0.083, *−*0.071]); UnDetect is ahead of XPert in every fold (top-1 0.258 to 0.287 against 0.186 to 0.201).

|  | top-1 |  |  |  |  | normalized rank |
|---|---|---|---|---|---|---|
| model | *n* = 2 | 3–5 | 6–10 | ≥ 11 | all | all |
| chance | 0.500 | 0.266 | 0.132 | 0.047 | 0.114 | 0.500 |
| CIGER | 0.627 | 0.386 | 0.223 | 0.099 | 0.185 | 0.412 |
| MultiDCP | 0.529 | 0.310 | 0.158 | 0.059 | 0.134 | 0.464 |
| DeepCE | 0.568 | 0.318 | 0.163 | 0.068 | 0.143 | 0.454 |
| TranSiGen | 0.661 | 0.393 | 0.229 | 0.118 | 0.202 | 0.397 |
| XPert | 0.645 | 0.375 | 0.218 | 0.110 | 0.191 | 0.404 |
| chemistry kernel | 0.678 | 0.438 | 0.270 | 0.145 | 0.233 | 0.368 |
| UnDetect | **0.754** | **0.506** | **0.334** | **0.169** | **0.271** | **0.326** |
| UnDetect + XPert | 0.727 | 0.468 | 0.309 | 0.154 | 0.252 | 0.342 |
| UnDetect − XPert | +0.108 | +0.132 | +0.116 | +0.059 | +0.080 | −0.077 |

### **A.13** Null tests for the knowledge views

Each view is tested against a null that keeps its coverage and marginal structure: target annotations permuted among the compounds that have them, and the one-hop STRING neighborhood of each compound’s targets taken from a degree-preserving rewired network [20] instead of the real one (table 20). The model is the line-specific term on LINCS P2-24 with one kernel per view.

**Table 20:** View null tests on LINCS P2-24 (7 lines): line-macro difference in Pearson, 95% bootstrap interval over the split’s compound groups, and lines with a positive difference. Targets and STRING: compounds with annot<u>ated targets; Cell Painting: compounds with a profil</u>e.

| contrast | plate-map split | ik14 split |
|---|---|---|
| targets − permuted targets | +0.035 [0.029, 0.042], 7/7 | +0.045 [0.037, 0.053], 7/7 |
| STRING one-hop − rewired | +0.000 [−0.003, 0.003], 5/7 | +0.001 [−0.001, 0.003], 6/7 |
| adding Cell Painting | +0.052 [0.045, 0.060], 7/7 | +0.088 [0.081, 0.095], 7/7 |

### **A.14** Data sources

All inputs are public (table 21); checksums of every downloaded file are kept with the code. Target and MOA annotations are literature-curated (Drug Repurposing Hub) and were not derived from L1000 profiles.

**Table 21:** Data sources. Checksums: MD5 or SHA-256 of the downloaded file, first eight hex digits.

| data | source and version | checksum |
|---|---|---|
| drug-blind benchmark | XPert data release [12] (Zenodo 17182939), as used in [3] | MD5 4a58bc01 |
| published predictions | Zenodo 22095049 (v2.0.0) [3] | SHA-256 manifest |
| LINCS L1000, Level 5 | GEO GSE92742 (Phase I), GSE70138 (Phase II) | GEO SHA-512 |
| DeepCE benchmark | pth1993/DeepCE, commit 9b0d04f | – |
| Tahoe-100M | tahoebio/tahoe-de-rhaister, rev. c7963cf;<br>pseudobulk deltas, fixed 2,000-gene panel | – |
| targets, MOA | CLUE LINCS2020 compoundinfo_beta; Tahoe drug table | SHA-256 a71fca6d |
| Cell Painting | broadinstitute/lincs-cell-painting, A549 48 h,<br>spherized consensus profiles (2026-10-03) | SHA-256 1e04d0be |
| protein interactions | STRING v12.0, human physical links | – |

## References

1. Ahlmann-Eltze, C., Huber, W., Anders, S.: Deep-learning-based gene perturbation effect prediction does not yet outperform simple linear baselines. Nature Methods 22(8), 1657–1661 (2025). 10.1038/s41592-025-02772-6

2. Álvarez, M.A., Rosasco, L., Lawrence, N.D.: Kernels for vector-valued functions: a review. Foundations and Trends in Machine Learning 4(3), 195–266 (2012). 10.1561/2200000036

3. Bai, J., Prince, S., Nitschke, G.S.: Deep learning models for chemical perturbation prediction do not yet utilise drug molecular features. bioRxiv (2026). 10.64898/2026.05.13.724458

4. Bray, M.A., Singh, S., Han, H., et al.: Cell Painting, a high-content image-based assay for morphological profiling using multiplexed fluorescent dyes. Nature Protocols 11(9), 1757–1774 (2016). 10.1038/nprot.2016.105

5. Cichonska, A., Pahikkala, T., Szedmak, S., Julkunen, H., Airola, A., Heinonen, M., Aittokallio, T., Rousu, J.: Learning with multiple pairwise kernels for drug bioactivity prediction. Bioinformatics 34(13), i509–i518 (2018). 10.1093/bioinformatics/bty277

6. Corsello, S.M., Bittker, J.A., Liu, Z., et al.: The Drug Repurposing Hub: a next-generation drug library and information resource. Nature Medicine 23(4), 405–408 (2017). 10.1038/nm.4306

7. Cortes, C., Mohri, M., Rostamizadeh, A.: Algorithms for learning kernels based on centered alignment. Journal of Machine Learning Research 13, 795–828 (2012)

8. Costello, J.C., Heiser, L.M., Georgii, E., et al.: A community effort to assess and improve drug sensitivity prediction algorithms. Nature Biotechnology 32(12), 1202–1212 (2014). 10.1038/nbt.2877

9. Evans, N.J., Mills, G.B., Wu, G., Song, X., McWeeney, S.: Graph structured neural networks for perturbation biology. bioRxiv (2024). 10.1101/2024.02.28.582164

10. Feng, J., Zhao, Z., Zhang, X., et al.: A knowledge-driven framework for predicting single-cell responses for unprofiled drugs. Nature Machine Intelligence 8(9), 1478–1491 (2026). 10.1038/s42256-026-01286-w

11. Gönen, M., Margolin, A.A.: Drug susceptibility prediction against a panel of drugs using kernelized Bayesian multitask learning. Bioinformatics 30(17), i556–i563 (2014). 10.1093/bioinformatics/btu464

12. Guo, Y., Zhang, H., Hu, H., Wu, J., Cao, J., Hsieh, C.Y., Yang, B.: Modelling drug-induced cellular perturbation responses with a biologically informed dual-branch transformer. Nature Machine Intelligence 8(1), 96–112 (2026). 10.1038/s42256-025-01165-w

13. Haghighi, M., Caicedo, J.C., Cimini, B.A., Carpenter, A.E., Singh, S.: High-dimensional gene expression and morphology profiles of cells across 28,000 genetic and chemical perturbations. Nature Methods 19(12), 1550–1557 (2022). 10.1038/s41592-022-01667-0

14. Hastie, T., Tibshirani, R.: Varying-coefficient models. Journal of the Royal Statistical Society: Series B (Methodological) 55(4), 757–779 (1993). 10.1111/j.2517-6161.1993.tb01939.x

15. Hetzel, L., Böhm, S., Kilbertus, N., Günnemann, S., Lotfollahi, M., Theis, F.: Predicting cellular responses to novel drug perturbations at a single-cell resolution. In: Advances in Neural Information Processing Systems 35. pp. 26711–26722 (2022)

16. Hodos, R., Zhang, P., Lee, H.C., et al.: Cell-specific prediction and application of drug-induced gene expression profiles. Pacific Symposium on Biocomputing 23, 32–43 (2018). 10.1142/9789813235533_0004

17. Hu, Q., Chen, Z., Gu, J.: Predicting drug-perturbed transcriptional responses using multi-conditional diffusion transformer. Quantitative Biology 14(1), e70016 (2026). 10.1002/qub2.70016

18. Kaufmann, L., Le Breton, L., Carraz-Billat, E., Fournier, Q., Lemieux, S.: A biological-response compound representation allows chemical perturbation prediction across cell lines. bioRxiv (2026). 10.64898/2026.09.28.755146

19. Lotfollahi, M., Klimovskaia Susmelj, A., De Donno, C., et al.: Predicting cellular responses to complex perturbations in high-throughput screens. Molecular Systems Biology 19(6), e11517 (2023). 10.15252/msb.202211517

20. Maslov, S., Sneppen, K.: Specificity and stability in topology of protein networks. Science 296(5569), 910–913 (2002). 10.1126/science.1065103

21. Molina, A., Zhang, X.: Perturbation response decomposition enables biologically aligned generalization to unseen perturbations and cellular contexts. bioRxiv (2026). 10.64898/2026.07.24.740459

22. Perez, E., Strub, F., de Vries, H., Dumoulin, V., Courville, A.: FiLM: Visual reasoning with a general conditioning layer. In: Proceedings of the AAAI Conference on Artificial Intelligence. vol. 32 (2018). 10.1609/aaai.v32i1.11671

23. Pham, T.H., Qiu, Y., Liu, J., Zimmer, S., O’Neill, E., Xie, L., Zhang, P.: Chemical-induced gene expression ranking and its application to pancreatic cancer drug repurposing. Patterns 3(4), 100441 (2022). 10.1016/j.patter.2022.100441

24. Pham, T.H., Qiu, Y., Zeng, J., Xie, L., Zhang, P.: A deep learning framework for high-throughput mechanism-driven phenotype compound screening and its application to COVID-19 drug repurposing. Nature Machine Intelligence 3(3), 247–257 (2021). 10.1038/s42256-020-00285-9

25. Qi, X., Zhao, L., Tian, C., et al.: Predicting transcriptional responses to novel chemical perturbations using deep generative model for drug discovery. Nature Communications 15, 9256 (2024). 10.1038/ s41467-024-53457-1

26. Rogers, D., Hahn, M.: Extended-connectivity fingerprints. Journal of Chemical Information and Modeling **50**(5), 742–754 (2010). 10.1021/ci100050t

27. Roohani, Y., Huang, K., Leskovec, J.: Predicting transcriptional outcomes of novel multigene perturbations with GEARS. Nature Biotechnology 42(6), 927–935 (2024). 10.1038/s41587-023-01905-6

28. Stock, M., Pahikkala, T., Airola, A., De Baets, B., Waegeman, W.: A comparative study of pairwise learning methods based on kernel ridge regression. Neural Computation 30(8), 2245–2283 (2018). 10.1162/neco_a_01096

29. Subramanian, A., Narayan, R., Corsello, S.M., et al.: A next generation connectivity map: L1000 platform and the first 1,000,000 profiles. Cell 171(6), 1437–1452 (2017). 10.1016/j.cell.2017.10.049

30. Svensson, V., Khan, U., Heydari, H., et al.: Back to basics: Observed statistics are sufficient to predict drug responses. bioRxiv (2026). 10.64898/2026.06.09.731197

31. Szklarczyk, D., Kirsch, R., Koutrouli, M., et al.: The STRING database in 2023: protein-protein association networks and functional enrichment analyses for any sequenced genome of interest. Nucleic Acids Research 51(D1), D638–D646 (2023). 10.1093/nar/gkac1000

32. Tong, X., Qu, N., Kong, X., et al.: Deep representation learning of chemical-induced transcriptional profile for phenotype-based drug discovery. Nature Communications 15, 5378 (2024). 10.1038/ s41467-024-49620-3

33. Viñas Torné, R., Wiatrak, M., Piran, Z., Fan, S., Jiang, L., Teichmann, S.A., Nitzan, M., Brbić, M.: Systema: a framework for evaluating genetic perturbation response prediction beyond systematic variation. Nature Biotechnology 44(6), 1050–1059 (2026). 10.1038/s41587-025-02777-8

34. Way, G.P., Natoli, T., Adeboye, A., et al.: Morphology and gene expression profiling provide complementary information for mapping cell state. Cell Systems 13(11), 911–923 (2022). 10.1016/j.cels.2022.10.001

35. Wong, D.R., Hill, A.S., Moccia, R.: Simple controls exceed best deep learning algorithms and reveal foundation model effectiveness for predicting genetic perturbations. Bioinformatics 41(6), btaf317 (2025). 10.1093/bioinformatics/btaf317

36. Wu, Y., Liu, Q., Qiu, Y., Xie, L.: Deep learning prediction of chemical-induced dose-dependent and context-specific multiplex phenotype responses and its application to personalized Alzheimer’s disease drug repurposing. PLoS Computational Biology 18(8), e1010367 (2022). 10.1371/journal.pcbi.1010367

37. Zhang, J., Ubas, A.A., de Borja, R., et al.: Tahoe-100M: A giga-scale single-cell perturbation atlas for context-dependent gene function and cellular modeling. bioRxiv (2025). 10.1101/2025.02.20.639398

